# PID-controller enhanced artificial *β*-cells

**DOI:** 10.1101/2025.01.27.634987

**Authors:** Lin Liu, Bruna Jacobson, Darko Stefanovic

## Abstract

Conventional management of diabetes via injection or external insulin pumps suffers from inconvenience and inability to accurately maintain blood glucose levels. A potential solution to these problems consists of implanting synthetic artificial *β*-cells that can sense glucose and transcribe insulin protein. We focus on one such artificial *β*-cell recently described in the work of Xie et al. Experimental results show these cells are able to release insulin and somewhat improve postprandial glucose levels in diabetic mice. However, they fail to achieve the degree of glucose regulation as in healthy mice. In our analysis, we explain that this artificial *β*-cell system has a major disadvantage: it is a high-dimensional dynamic system but with little tuning space. Here, we propose a PID-controller-based enhanced artificial *β*-cell design to solve this issue. Our results based on an analytical model and numerical simulations show that the computational method of PID-control can enhance engineered artificial *β*-cells, such that they could perform better in regulating glucose levels in Type 1 diabetic mice compared with artificial *β*-cells without PID-control: they could shut down the production of insulin in time and maintain a proper glycemia level, and there is more tuning space for artificial *β*-cells with PID control.

## Introduction

Diabetes is a chronic illness that affects the body’s ability to convert food into energy.^1^ Many mammalian species are susceptible to diabetes, including humans, mice, pigs, and dogs.^2^ In a healthy body, carbohydrates are converted into glucose, which provides energy for daily activities. In response to an increase in blood glucose, the *β*-cells in the pancreas release insulin. Insulin in turn allows blood glucose to enter cells so that it can be used as fuel. In diabetic animals, insulin production is insufficient or insulin is not utilized as efficiently as in healthy animals because the cells in muscles, fat, and liver do not respond to it. In either case, high blood glucose levels occur, which can lead to serious health problems, including heart disease, kidney disease, stroke, and lower limb amputations.^3^ There are several phenotypes of diabetes, and the two most common are Type 1 diabetes (T1D) and Type 2 diabetes (T2D). While T2D involves insulin resistance, owing to aging or obesity, T1D is characterized by the absence of functional *β*-cells from birth in animals and humans. There is currently no prevention for T1D. Patients with T1D must take in external insulin every day to maintain their blood glucose level in the normal range. Traditional treatment for diabetes includes oral medication intake, insulin injections, and insulin pumps, which all suffer from inconvenience, inaccuracy, and hygiene issues such as risk of skin infection.^4^ In 2016, Xie et al. proposed a novel approach for diabetic treatment, the artificial *β*-cell.^5^ In particular, they introduce HEK-*β* cells, which are genetically modified from the monoclonal HEK-293 cell line. According to this design, when there is a high concentration of glucose in the cell exterior environment, the ATP concentration in the cell increases and subsequently ion channels are supposed to be triggered. The final result of the ion channel activity is the transcription of mRNA and the subsequent production and secretion of insulin to lower the blood glucose levels. Xie et al. also offer a detailed analytical model for these cells in a mouse model. Here we refer to these cells as artificial *β*-cells for short. We can view the ion influx as a biological sensor of glucose, and the ATP as a real-time indicator of glucose levels. However, while these cells provide a degree of insulin regulation when implanted in a diabetic mouse, the regulation is not comparable to that of a healthy mouse. Our analysis shows that a plausible explanation lies in the low number of artificial *β*-cells implanted. However, if the number of artificial *β*-cells is increased, we find that a very high blood insulin level persists that is lethal to mice. A detailed analysis of the model indicated that this is due to an insufficiently sensitive signaling pathway from ATP to mRNA: the insulin production does not cease on time, even when the external glucose levels and the internal ATP levels are dangerously low.

Having examined these limitations in the achieved performance of artificial *β*-cells, we explore, numerically, what might be achievable if the section of the signaling pathway from ATP to mRNA were replaced by an explicit control circuit. Here we do not fully specify the design of that circuit, leaving it to a future paper, but we note that any such design will have to augment or modulate the machinery of the ion channels present in the cell. In choosing which control algorithm to realize first, a natural choice is simple feedback control, namely proportional-integral-derivative (PID) control, which has been used in external insulin pump control.^6–8^ Therefore, we introduce a model of PID-controlled *β*-cells for glucose level regulation in diabetic patients.

PID control is also attractive because it has been shown that a chemical reaction network (CRN) with deterministic mass action semantics can implement integral, proportional, and derivative operations over chemical species concentrations, thereby implementing PID controller logic,^9–11^ wherein the integral, proportional, and derivative gain can be adjusted by appropriately setting the reaction rates. Here, we do not specify the components of such a CRN; instead we treat the PID controller abstractly and we focus on the feasibility of glucose level regulation with this class of controller, and the tuning of the PID gain parameters.

We show that the PID *β*-cells we propose can provide a more effective diabetic treatment. In the Methods and Results section, we report that PID *β*-cells can outperform the original artificial *β*-cells, as we show via computer simulation: Type 1 diabetic mice implanted with PID *β*-cells have postprandial glucose level time courses closer to those of healthy mice, compared with T1D mice implanted with the original artificial *β*-cells without PID control. Moreover, the PID *β*-cells can also provide more tuning space on the glucose time courses compared with artificial *β*-cells, by varying controller parameters. In the Discussion section we touch on the advantages and the limitations of PID *β*-cells.

## Background on artificial *β*-cells

Through biochemical reactions, the ion channels on the membrane of the artificial *β*-cells function as a sensing circuit that controls the expression of insulin-secreting genes. The cells can roughly be viewed as a feedback controller as shown in Figure 1: when there is a high concentration of glucose in the cell exterior environment, the ATP concentration in the cell also increases and subsequently ion channels are triggered. The final result of the ion channel activity is the transcription of mRNA and the subsequent production and secretion of insulin to lower the blood glucose levels. After the glucose level drops, the ATP level also decreases, and the ion channels are supposed to shut off insulin production. Three groups of mice were used to test the effectiveness of artificial *β*-cells, healthy wild type (WT) mice, mice with T1D without any treatment, and T1D mice with artificial *β*-cell implants. In oral glucose tolerance tests using sugared water, glucose levels were measured every 30 minutes up to 150 minutes and every 60 minutes afterwards (Figure 2). T1D mice implanted with artificial *β*-cells (blue dots) showed lower blood glucose levels compared with T1D mice (red dots), but not as low as healthy mice (black dots). It appears that the glucose level in T1D mice treated with artificial *β*-cells is decreasing too slowly, failing to reach the healthy glucose range (less than 5.5 mM)^12^ within two hours. In addition to these in vivo experiments, Xie et al. presented a unified mathematical model of the artificial *β*-cells (with details of the processes of glycolysis, ion channel trans-duction, transcription, translation, and secretion), the implant delivery capsules, and the mouse glucose metabolism for the in vitro experiments. They use that model to analyze and explain the oral glucose tolerance test results for each group of mice. We reproduced this model in Matlab, handled some vagueness, and corrected some possible errors (see Supplementary material). There are dozens of ordinary differential equations and variables in the model to describe the dynamics of blood glucose level and the artificial *β*-cells. Specifically, the model simulates the process of increasing ATP level in the artificial *β*-cell as a result of a high extracellular glucose level, as well as the subsequent mRNA transcription and insulin production as a result of ion channel activities for T1D mice implanted with artificial *β*-cells. The model is also capable of simulating long-lasting high glucose levels in T1D mice without any treatment as well as a changing glucose level pattern in healthy mice. Of main interest to us is the oral glucose intake test, corresponding to the experimental results which we reproduce in Figure 2. This is a three-stage protocol (cf. Ref. 5, Table S3, SI p. 40) as follows: Stage 1: set all variables to initial values with respect to Table 1 and Table 5 in the supplementary material, set the insulin production factor *F*_*I*_ to 0.001 for T1D mice and 1 for healthy mice, and simulate for 72 hours without the cells implant (by setting *γ*_*I*_ and *γ*_*G*_ to zero). At the end of Stage 1, we reach state *S*_11_ for healthy mice and state *S*_12_ for T1D mice. Simulating Stage 2 involves first setting *G*_*S*_ to 0 for *S*_11_ and *S*_12_ then following by three scenarios: (1) With *S*_11_ as initial condition, we simulate for 21 days for healthy mice without implant, and we reach state *S*_21_. (2) With *S*_12_ as initial conditions, we simulate for 21 days for T1D mice without implant, and we reach state *S*_22_. (3) With *S*_12_ as initial condition, we add the artificial *β*-cell implant to T1D mice, then simulate for 21 days to reach state *S*_23_. Here, the implant is added by setting *γ*_*I*_ and *γ*_*G*_ to positive values according to Table 8 in the supplementary material and setting cell density to 1.17 × 10^7^ cells/L. The protocol can be represented as in Figure 3. This last simulation serves the purpose of making the artificial *β*-cells “mature”: the inner states of the cell are balanced with the outer steady-state environment. Lastly, we have Stage 3, the actual oral glucose test. We manually set *G*_*I*_, the glucose level in the mouse stomach, to 63.16 mM in states *S*_21_, *S*_22_, and *S*_23_, respectively, then perform three simulations: (1) With *S*_21_ as initial condition, we simulate healthy mice without any implant for six hours; the result is the black curve in Figure 2. (2) With *S*_22_ as initial condition, we simulate T1D mice without any implant for six hours; the result is the red curve in Figure 2. (3) With *S*_23_ as initial condition, we simulate T1D mice with implant for six hours (cell density 1.17 × 10^7^ cells/L); the result is the blue curve in Figure 2. Focusing on Stage 3, after a certain amount of time, artificial *β*-cells implanted in T1D mice indeed successfully decrease glucose levels compared with T1D mice without treatment. But in general, the glucose curve of the T1D group does not follow the same pattern as the healthy group: it drops too slowly and does not reach normal range within six hours.

**Table 1:** The glucose and insulin related variables.

| Name | Definition | Unit | Initial value |
| --- | --- | --- | --- |
| $G$ | Concentration of glucose in the blood | mM | 0 |
| $G_I^{***}$ | Concentration of glucose in an additional virtual compartment | mM | 0 |
| $I_a$ | Concentration of insulin in an action compartment | $\mu\text{g/L}$ | 0 |
| $I_i$ | Concentration of insulin in a virtual compartment | $\mu\text{g/L}$ | 0 |
| $I$ | Concentration of insulin in the blood | $\mu\text{g/L}$ | 0 |

**Table 2:** The glucose and insulin related constants.

| Name | Definition | Value | Unit | Source |
| --- | --- | --- | --- | --- |
| $g_p$ | Glucose production | 0.1061 | mM/min | Estimated |
| $g_r$ | Insulin-independent glucose removal | $4.1 \times 10^{-3}$ | mM/min | Estimated |
| $k_{si}$ | Insulin sensitivity | $1.2 \times 10^{-3}$ | L/$\mu\text{g}$/min | Estimated |
| $V_S$ | Volume of the synthetic compartment | $1.5 \times 10^{-4}$ | L | Fixed |
| $V_B$ | Volume of the mouse blood | $3.35 \times 10^{-4}$ | L | Fixed |
| $k_{iar}$ | Insulin clearance from action compartment | 0.0566 | /min | Estimated |
| $k_{iir}$ | Insulin clearance from virtual compartment | 0.0351 | /min | Estimated |
| $F_I^{***}$ | Insulin production factor of T1D mice | 0.001 | - | Fixed |
| $F_I^{***}$ | Insulin production factor of healthy mice | 1 | - | Fixed |
| $k_p$ | Basal insulin production rate | $4.0338 \times 10^{-5}$ | $\mu\text{g/L/mM/min}$ | Estimated |
| $k_i$ | Insulin mediated insulin production rate | $1.05 \times 10^{-10}$ | $\mu\text{g/L/mM/min}$ | Estimated |
| $k_d$ | Glucose mediated insulin production rate | 0.234 | $\mu\text{g/L/mM/min}$ | Estimated |
| $K_{MI}^{***}$ | Base concentration of insulin | 0.01 | $\mu\text{g/L}$ | 17 |
| $n_{HI}^{***}$ | Hill coefficient | 8 | - | 17 |

**Table 3:**
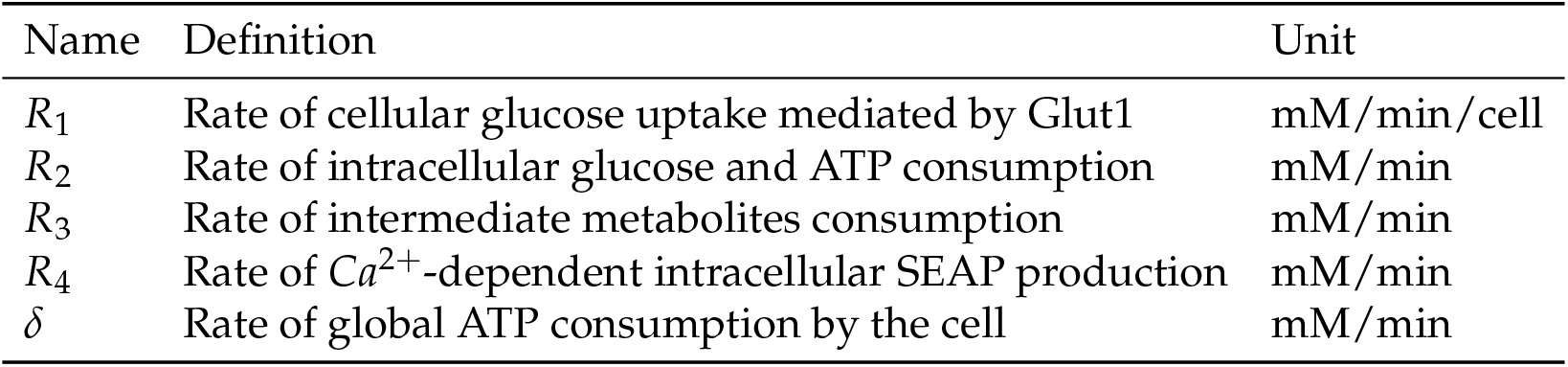
The rates related variables.

**Table 4:** The rates related constants.

| Name | Definition | Value | Unit | Source |
| --- | --- | --- | --- | --- |
| $q$ | Autocatalytic stoichiometry | 1 | - | Estimated |
| $V_{\max 1}$ | Maximum specific rate of cellular glucose uptake | $10^{-13}$ | mmol/min/cell | 18 |
| $K_{m1}$ | Michaelis-Menten constant for the uptake of glucose | 0.31 | mM | Estimated |
| $V_{\max 2}$ | Max rate of intracellular glucose and ATP consumption | 10 | mM/min | Estimated |
| $K_{m2A}$ | Affinity constant for allosteric inhibition of ATP consumption by ATP | 1.61 | mM | Estimated |
| $K_{m2G}$ | Michaelis-Menten constant for glucose consumption | 1.95 | mM | Estimated |
| $a$ | Hill coefficient for binding of ATP to phosphofructokinase | 1 | - | Estimated |
| $b$ | Hill coefficient for allosteric inhibition of ATP consumption by ATP | 1 | - | Estimated |
| $c$ | Hill coefficient for allosteric inhibition of intermediate metabolites consumption by ATP | 1 | - | Estimated |
| $V_{\max 3}$ | Max. intermediate consumption rate | 9.96 | mM/min | Estimated |
| $K_{m3}$ | Affinity for intermediary consumption inhibition by ATP | 1.83 | mM | Estimated |
| $A_{tot}$ | Total concentration of ATP + ADP | 5 | mM | 19 |
| $u$ | Max global rate of ATP consumption | 0.12 | mM/min | Estimated |
| $K_{\delta}$ | Michaelis-Menten constant for ATP consumption | 2 | mM | Estimated |
| $d_4^{***}$ | Hill coefficient for calcium-dependent gene expression | 1.6 | - | Estimated |

**Table 5:** The ion related variables.

| Name | Definition | Unit | Initial value |
| --- | --- | --- | --- |
| $N_S$ | Cell density in the synthetic compartment | mM | 0 |
| $G_S$ | Concentration of glucose in the synthetic compartment | mM | 0 |
| $G_{Si}$ | Concentration of glucose in the cell | mM | 0 |
| 6CP | Intermediate metabolites (phosphorylated six-carbon sugars) concentration in the cell | mM | 0 |
| $A$ | Concentration of ATP in the cells | mM | 0.1 |
| $Ca_S$ | Concentration of calcium ions in the synthetic compartment | mM | 1.8 |
| $Ca_{Si}$ | Concentration of calcium ions in the cells | mM | $2 \times 10^{-5}$ |
| $K_S$ | Concentration of potassium ions in the synthetic compartment | mM | 5.3 |
| $K_{Si}$ | Concentration of potassium ions in cells | mM | 200 |
| $Na_S$ | Concentration of sodium ions in the synthetic compartment | mM | 155 |
| $Na_{Si}$ | Concentration of sodium ions in cells | mM | 8 |
| $V$ | Transmembrane voltage $\phi_{in} - \phi_{out}$ | mV | -90 |
| $M_S$ | Concentration of insulin mRNA in the cells | mM | 0 |
| $I_S$ | Concentration of insulin in the synthetic compartment | $\mu\text{g/L}$ | 0 |

**Table 6:**
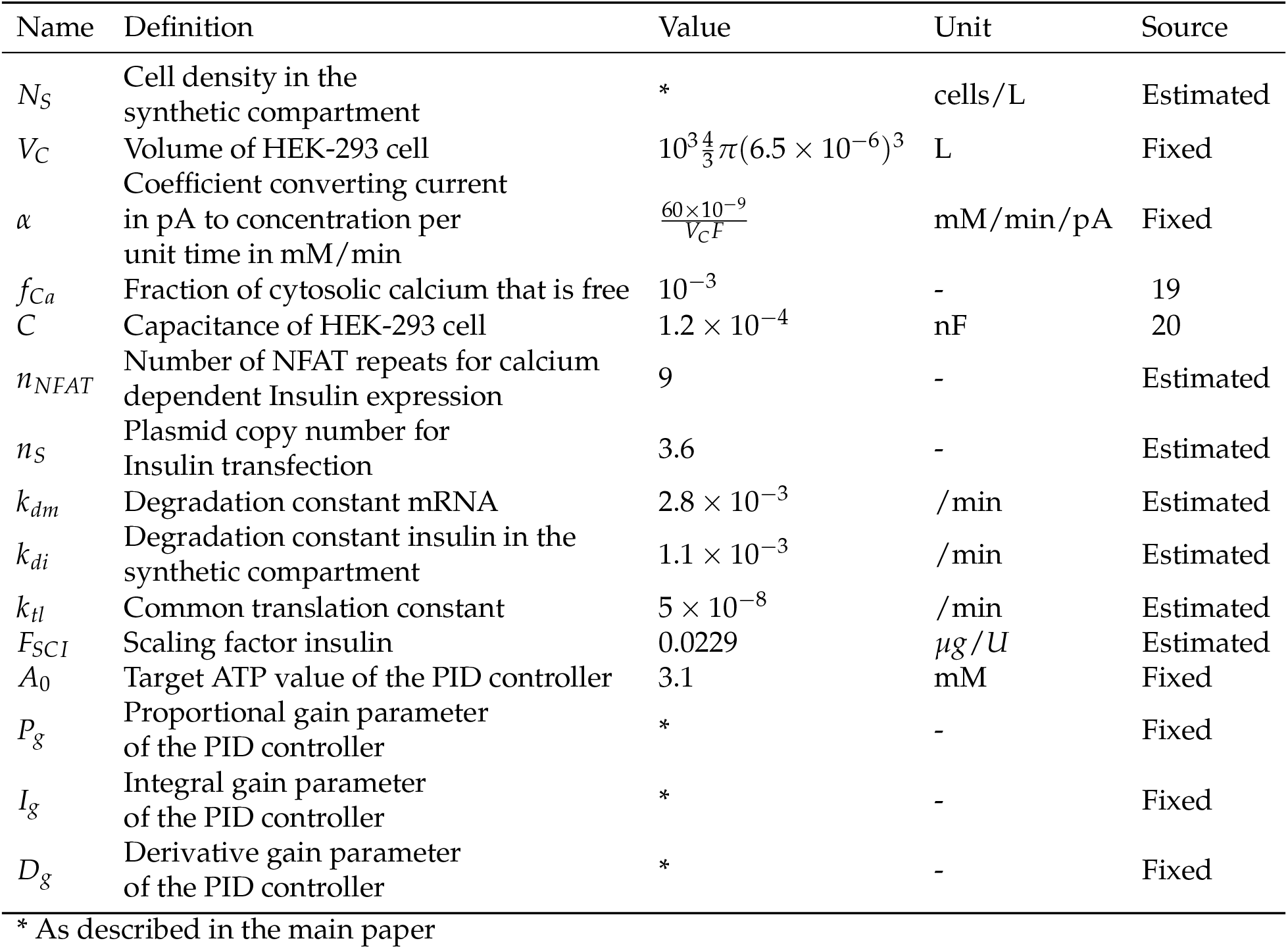
The rates related constants.

**Table 7:**
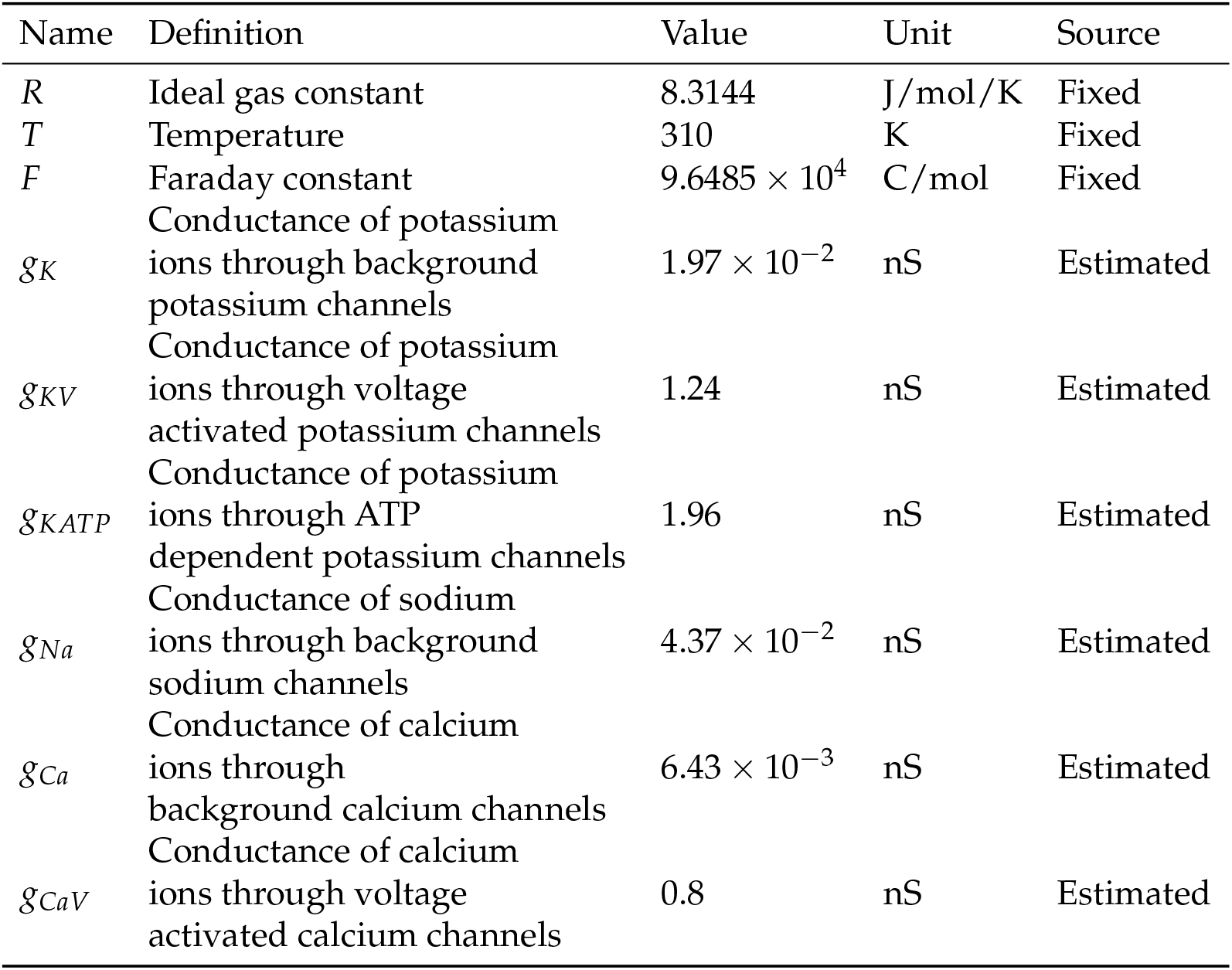
The flux transfer related constants.

| Name | Definition | Value | Unit | Source |
| --- | --- | --- | --- | --- |
| $R$ | Ideal gas constant | 8.3144 | J/mol/K | Fixed |
| $T$ | Temperature | 310 | K | Fixed |
| $F$ | Faraday constant | $9.6485 \times 10^4$ | C/mol | Fixed |
| $g_K$ | Conductance of potassium ions through background potassium channels | $1.97 \times 10^{-2}$ | nS | Estimated |
| $g_{KV}$ | Conductance of potassium ions through voltage activated potassium channels | 1.24 | nS | Estimated |
| $g_{KATP}$ | Conductance of potassium ions through ATP dependent potassium channels | 1.96 | nS | Estimated |
| $g_{Na}$ | Conductance of sodium ions through background sodium channels | $4.37 \times 10^{-2}$ | nS | Estimated |
| $g_{Ca}$ | Conductance of calcium ions through background calcium channels | $6.43 \times 10^{-3}$ | nS | Estimated |
| $g_{CaV}$ | Conductance of calcium ions through voltage activated calcium channels | 0.8 | nS | Estimated |

**Table 8:** The flux transfer related constants, continued.

| Name | Definition | Value | Unit | Source |
| --- | --- | --- | --- | --- |
| $n_C$ | plasmid copy number for Cav1.3 transfection | 3.6 | - | Estimated |
| $K_K$ | Rate constant of potassium uptake from the medium by NaK pump | 1 | /min | 21 |
| $I_{NaK}^{max}$ | Maximum sodium-potassium pump current | 42.72 | pA | Estimated |
| $I_{CaP}^{max}$ | Maximum calcium pump current | 9.4 | pA | Estimated |
| $\gamma_I$ | Vascular exchange constant insulin | 0.9137 | /min | Estimated |
| $\gamma_G$ | Vascular exchange constant glucose | 0.7074 | /min | Estimated |
| $D_G$ | Vascular exchange constant glucose | 0.2986 | /min | Estimated |
| $A_{tot}$ | Total concentration of ATP + ADP | 5 | mM | 19 |

**Figure 1:**
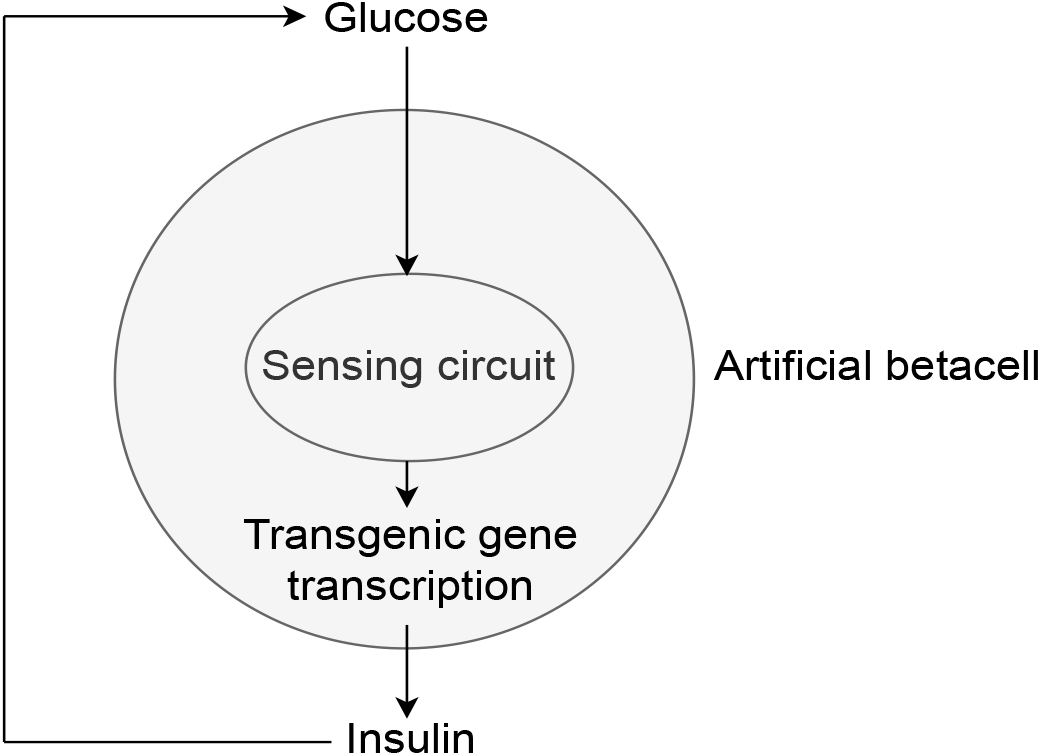
Model of the closed loop diagram of a HEK-293 transgenic artificial *β*-cell, from Xie et al. (2016). Glucose levels are sensed by ion channels on the cell membrane (sensing circuits), which trigger the transcription of the transgenic insulin-producing gene and the subsequent secretion of insulin, lowering glucose blood levels.

**Figure 2:**
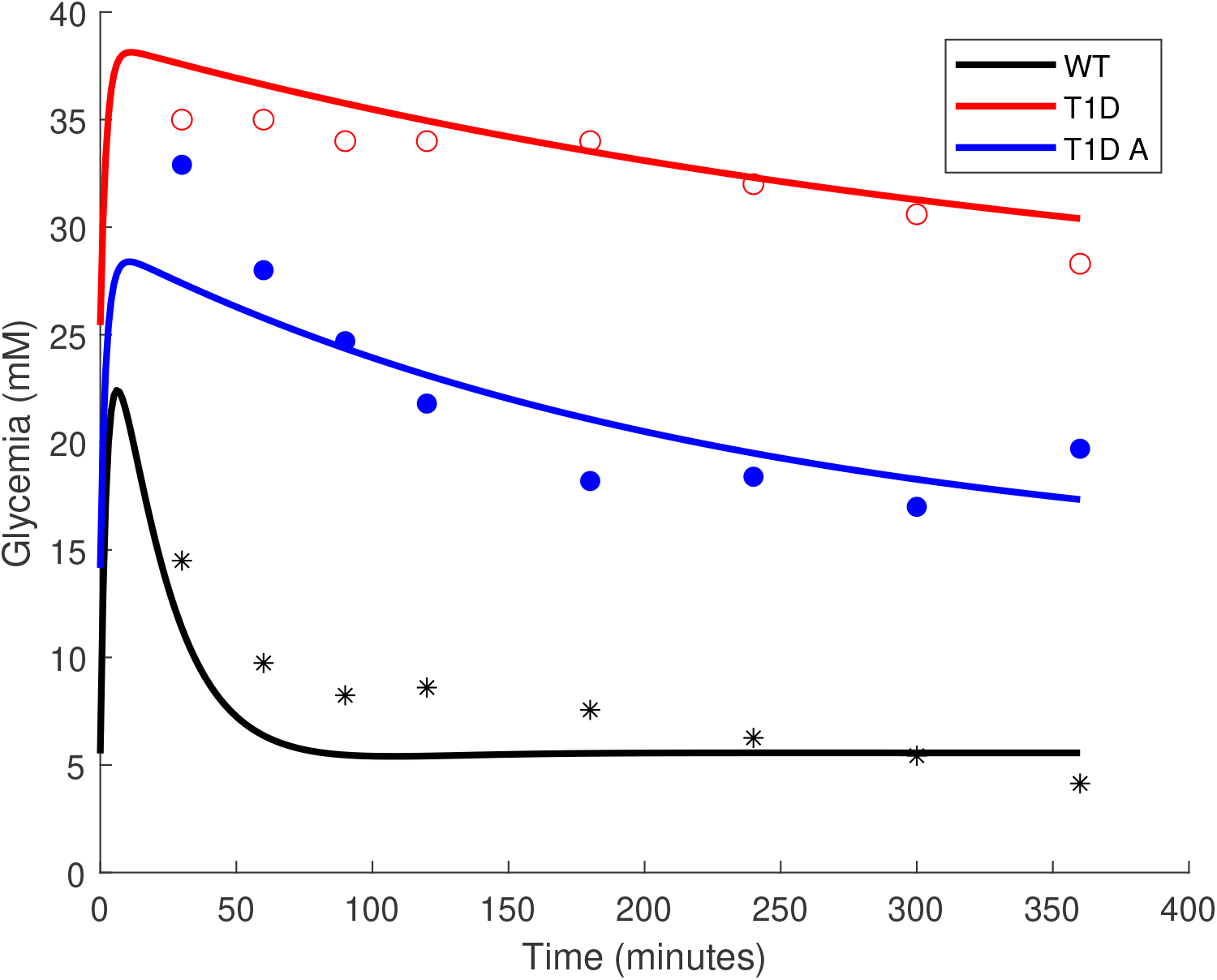
The dots represent the mean blood glucose levels of different mice groups in a six-hour time course following an oral glucose test (as read from Xie et al. (2016), Figure 4(G)). Before the oral glucose test, the mice had been fasting for an extended period of time. In the experiment, artificial *β*-cells are placed inside capsules. By diffusion, the capsules exchange insulin or glucose with the blood of the T1D mice. Red hollow dots represent the mean recorded experimental values for T1D mice without any treatment; black stars represent healthy mice; blue solid dots represent T1D mice treated with artificial *β* cells. The solid lines are our simulation results as the reconstruction of Xie et al. (2016), Figure 4(G). “WT” is the healthy mice group represented by the black curve, “T1D” is the T1D mice group without any treatment represented by the red curve, and “T1D A” is the T1D mice group treated with artificial *β*-cells represented by the blue curve. The cell density is set to 1.17 × 10^7^ cells/L to be visually consistent with experiment results; other parameters are as reported. It is evident that the artificial *β* cells treatment has some effect on the T1D mice, however the rate of glycemia decay is not as fast as in healthy mice.

**Figure 3:**
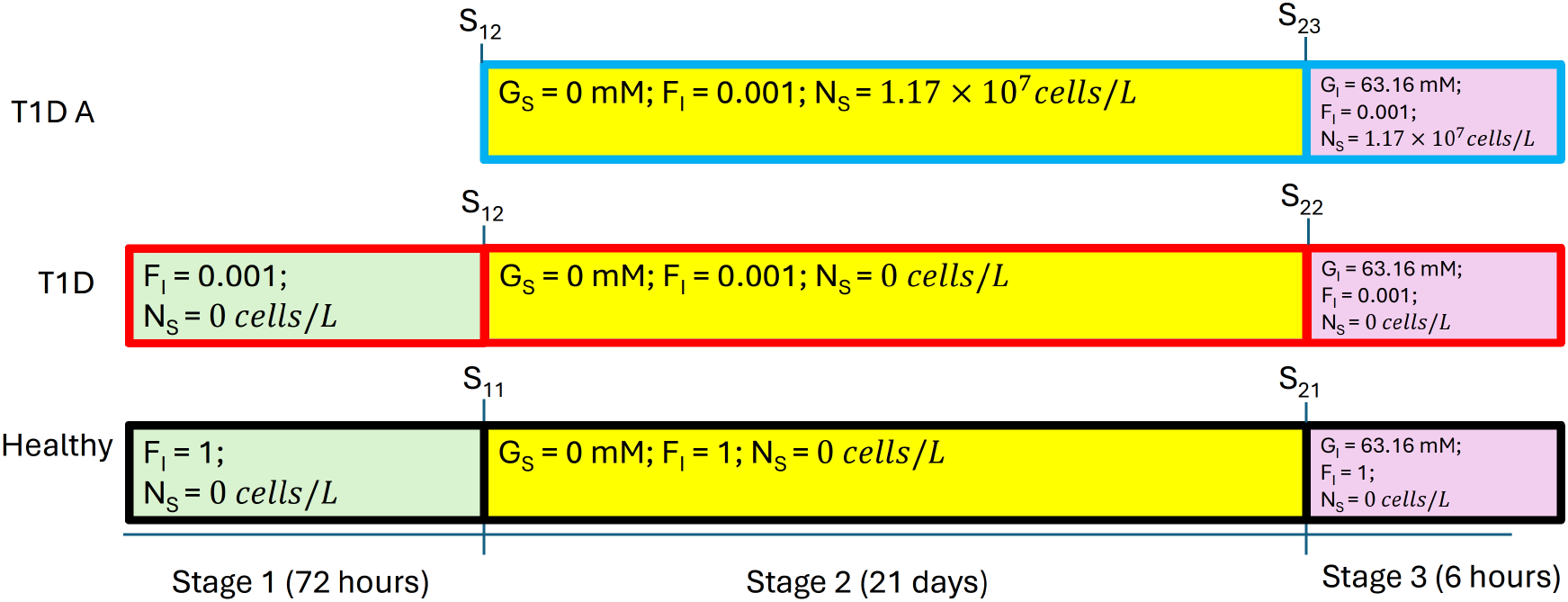
The protocol used for the oral glucose test. The initial values used in Stage 1 can be found in Table 1 and Table 5 of the supplementary material.

## Methods and results

From Figure 2, it can be seen that the glucose level in T1D mice implanted with artificial *β*-cells (of density value 1.17 × 10^7^ cells/L) is always too high after oral glucose intake, compared with healthy mice. There can be two possible causes: either there is insufficient quantity of artificial *β*-cells implanted, or the quantity is sufficient but the cells are not functioning at their peak level for producing insulin. To figure out which is the most likely cause, we plotted the mRNA concentration in the cell over the six-hour time course of Stage 3 in Figure 4. It shows that the mRNA concentration in the cells reaches 0.03087 mM quickly and remains at that value, which is very close to the upper limit of intracellular mRNA concentrations.^13^ Therefore, we can assume that the cells are functioning at their peak level.

**Figure 4:**
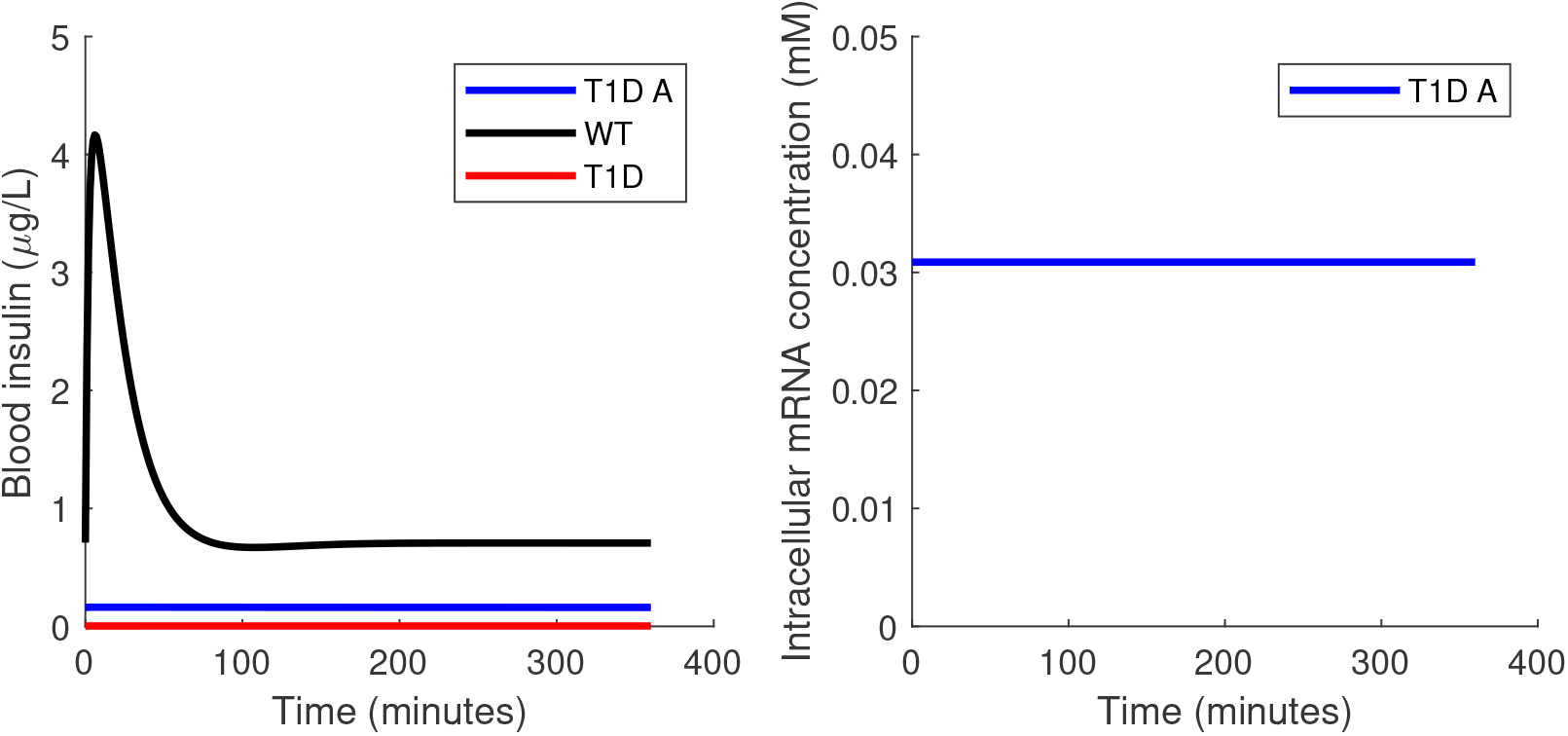
The insulin concentration in blood (left panel) and the mRNA concentration (right panel) in cells during Stage 3 (the six-hour time course after the oral glucose test). The implanted *β*-cell density is 1.17 × 10^7^ cells/L. The legend is the same as for Figure 2. Notice that in the right panel there is only one curve since the intracellular mRNA concentration is only meaningful in the T1D mice group with implanted cells. The artificial *β* cells are working at their full capacity as evidenced by the high level of mRNA.

We now consider insufficient quantity of cells as a possible cause of imperfect control. To increase the quantity of cells, we can either increase the cell density or the volume. As the volume is fixed by the implantation procedure, here we vary the cell density alone. Since the natural *β*-cell density is around 10^12^ cells/L for healthy mice^14^ and the artificial *β*-cells are of similar size,^13^ we increased the density to 5 × 10^11^ cells/L. This change, however, causes a problem during the second stage of the protocol: the glucose drops to an extremely low level (0.238 mM) within 20 minutes of the second stage. This is a dangerously low glucose level for the mice.^14^ Therefore, if we implant a high density value of the *β*-cells that is close to the healthy mice in the second stage, the mice are unlikely to survive during this stage. Hence, we changed the protocol to the following: Stage 1: same as stage 1 in the description of the artificial *β*-cell protocol above; Stage 2: the same setup for the mice without implant, but we also implant a very low density (1 cell/L) to the T1D mice such that these cells with extremely low density do not have any practical effect on the mice. After three weeks of fasting, the internal states of the artificial *β*-cells are balanced with the outer environment of the T1D mice; Stage 3: everything is the same as stage 3 in the artificial *β*-cell background session except we instantaneously increase the implanted cell density value to 5 × 10^11^ cells/L. This is to simulate an oral glucose test right after the implant of high density mature *β*-cells as the treatment for T1D mice. The protocol for increased cell density value can be represented in Figure 5.

**Figure 5:**
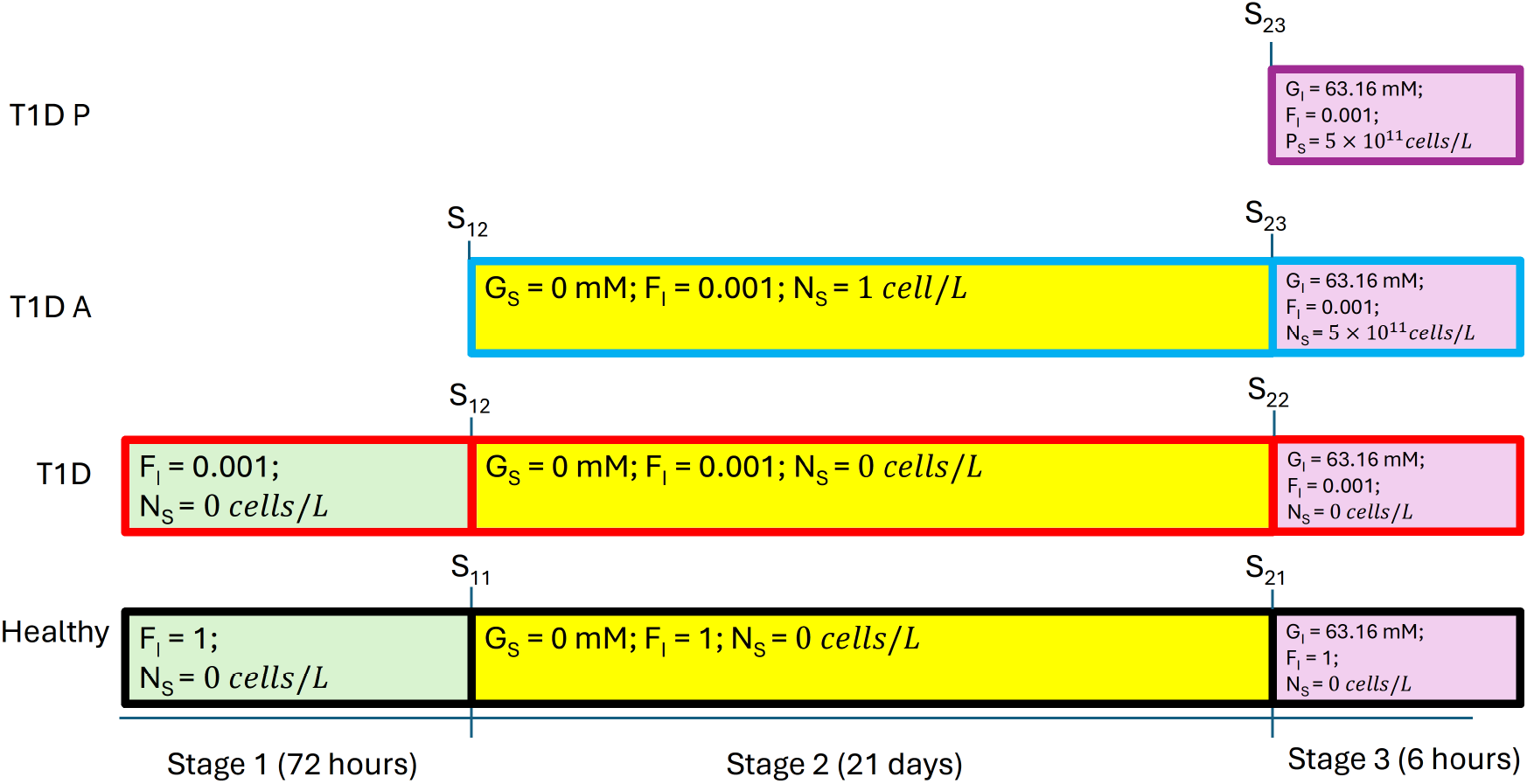
The protocol used with the cell density increased to 5 × 10^11^ cells/L. Note that *P*_*S*_ in Stage 3 means the PID *β*-cell density.

A plot of the results of the above-stated Stage 3 is shown in Figure 6. After increasing the cell density to 5 × 10^11^ cells/L, the blood insulin level can roughly reach 4 *µ*g/L within five minutes, which is comparable to that of the healthy mice. However, as a result of this increase in cell density, by 50 minutes the blood glucose level drops to near zero, which again is lethal. According to the simulation, we observe that the cells are still producing insulin almost to full capacity even after the glucose concentration has reached a healthy level from Figure 6. Therefore, it is necessary to introduce some mechanism that shuts off insulin production when necessary, and this is precisely what a PID controller can do.

**Figure 6:**
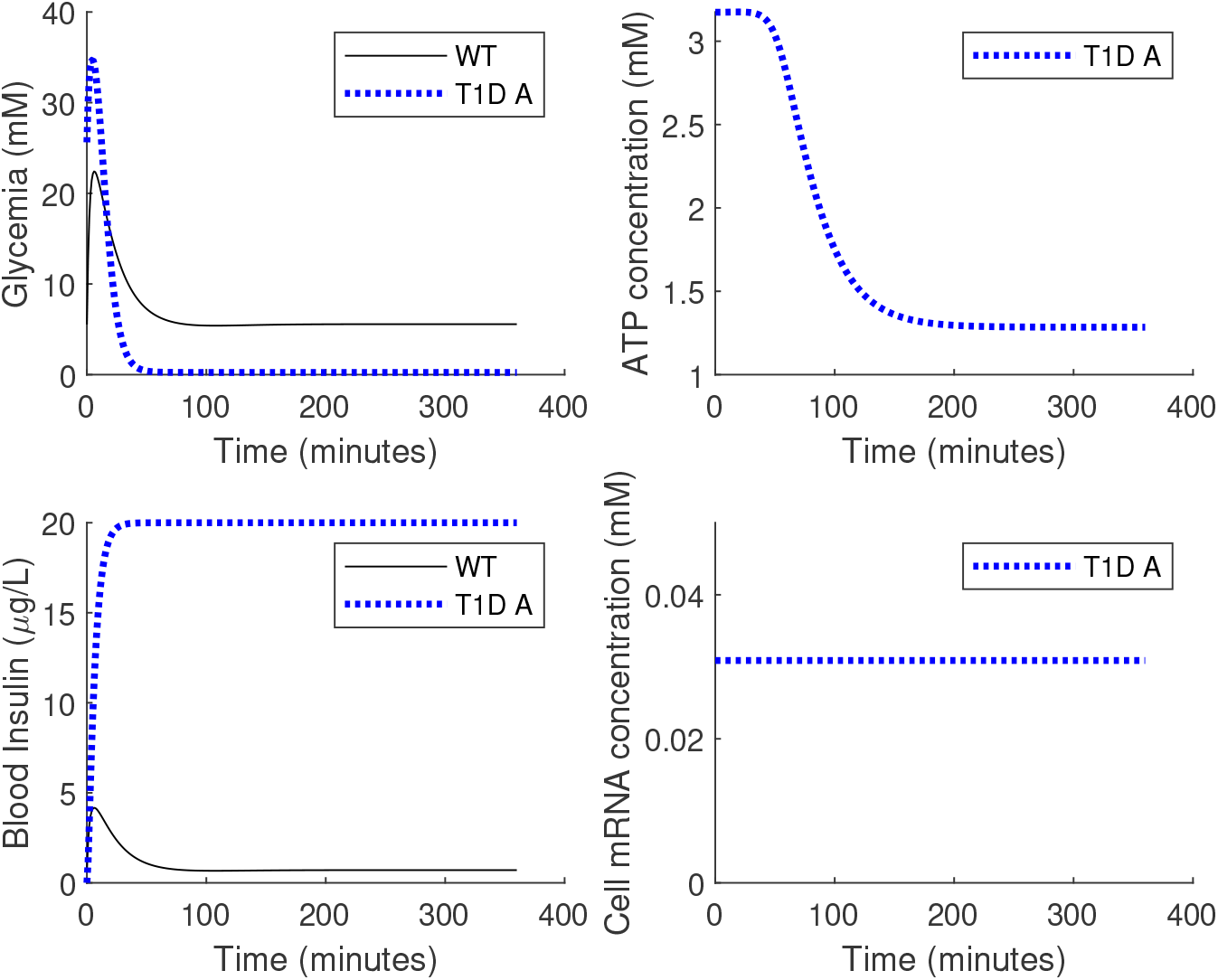
Setting the implanted artificial *β*-cell density to 5 × 10^11^ cells/L, the upper bound of blood insulin level is set to 20 *µ*g/L, consistent with highest possible mouse blood insulin level.^13^ The legend is the same as in Figure 2. Blue dotted line indicates there is a higher implant density. In response to the rapid drop of glucose levels when the cell density increases (upper left panel), the ATP also drops to a very low level (upper right panel). However, the artificial *β*-cells are unable to stop the production of insulin on time (lower left panel). The mRNA remains at a high level even when the ATP level is low due to the insensitivity of ion channels (lower right panel).

Our aim is to explore various control algorithms that can be implemented in the artificial *β*-cell. Due to CRN’s ability to implement the PID controller and our need to maintain the glucose level within a proper range, we selected the PID control and designed the PID *β*-cell. The following differential equation describes the PID controller as a conceptual replacement for the ATP-to-mRNA pathway in the PID *β*-cell:

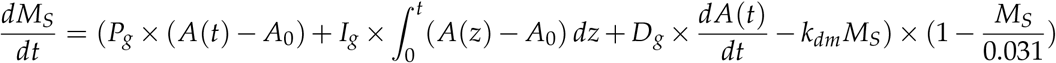

Here, A(t) is the concentration of ATP in the cells; *P*_*g*_, *I*_*g*_, and *D*_*g*_ represent the P, I, and D gain parameters respectively; *A*_0_ is the target value for ATP and is set to 3.2 mM. (Rounding 3.1737 mM, the ATP concentration after a long fasting period for T1D mice implanted with artificial *β*-cells with negligible density.)

We must also optimize the gain parameters. We use a constrained sampling approach, the constraint being that the blood glucose level must never drop below 4.14 mM (the mean blood glucose level in healthy mice after six hours of fasting, from in vivo experiments). We explore *P*_*g*_, *I*_*g*_, and *D*_*g*_ from 0 to 15 with a step size of 1. We limited the gain parameters to no more than 15 since in an eventual implementation they are determined by reaction rates from the CRN and we wish to avoid large ranges of required rates. Subject to the constraint, we select the glucose curve with the minimum *L*^2^ distance from the glucose curve of healthy mice. This is achieved with *P*_*g*_ = 14, *I*_*g*_ = 11, and *D*_*g*_ = 6; the results are shown in Figure 7.

**Figure 7:**
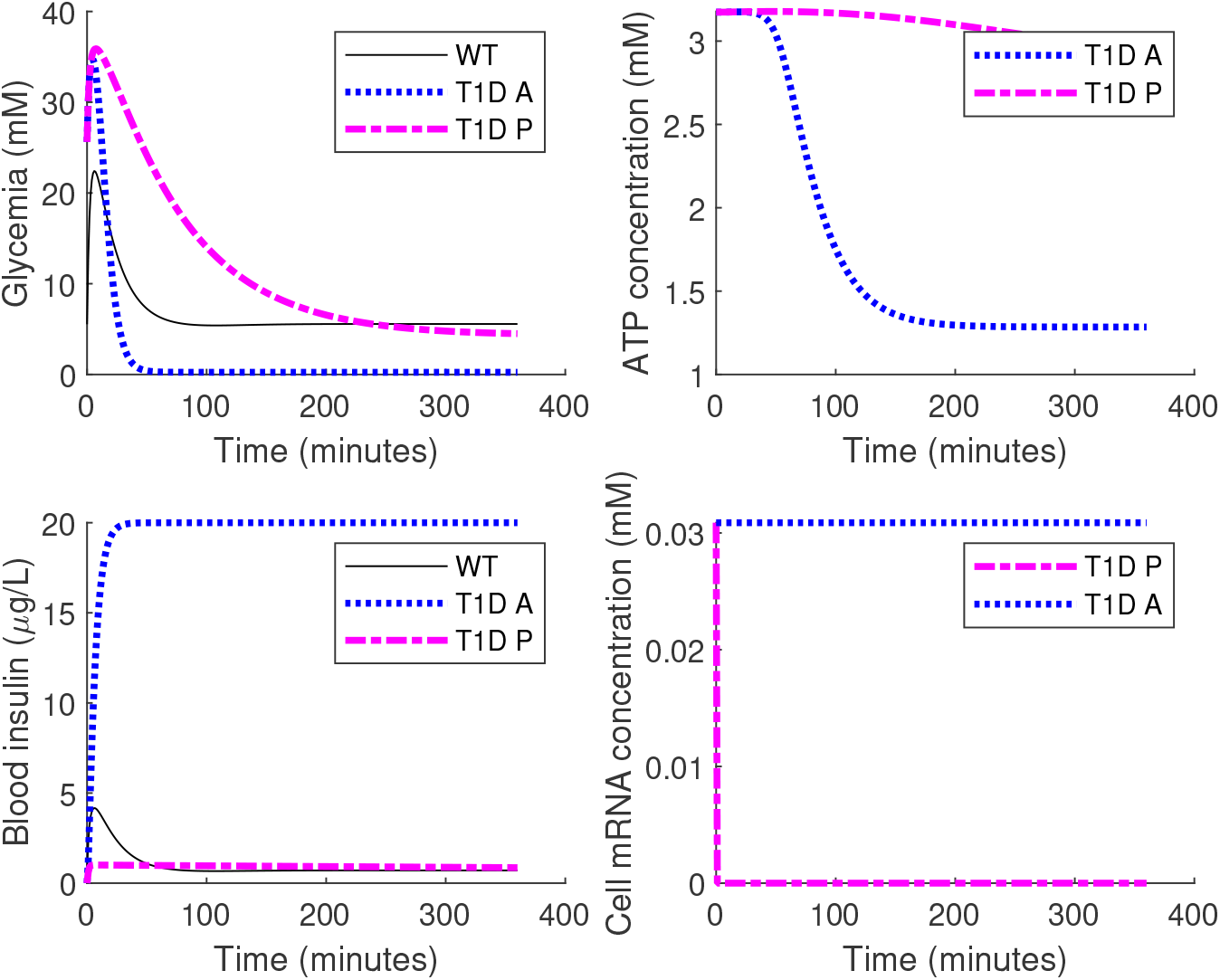
A comparison of artificial *β*-cells and PID *β*-cells. “T1D P” means the T1D mice treated with PID *β*-cells. The magenta dashed line represents the simulation for treatment result of T1D mice with PID *β*-cell implanted. The implanted PID *β*-cell density is the same as that of artificial *β*-cell, which is 5 × 10^11^ cells/L. For the PID controls, we have selected the gain parameters as *P*_*g*_ = 14, *I*_*g*_ = 11, and *D*_*g*_ = 6. Other legends are the same as Figure 2. For T1D mice treated with PID *β*-cells, the glucose level converges to a normal level (upper left panel), as does the ATP level (upper right panel). For mice treated with PID *β*-cell, the blood insulin level does not explode (lower left panel) since mRNA, the switch for insulin production, is shut off on time (lower right panel).

After 110 minutes, the blood glucose level of T1D mice implanted with PID *β*-cells drops below 10 mM and then remains around 5 mM throughout the experiment. The mRNA concentration in PID *β*-cells drops to zero after the first minute, and the secretion of insulin stops, as desired. Thus both hyperglycemia and hypoglycemia are avoided.

The PID *β*-cell also provides a more flexible tuning space compared with the artificial *β*-cell: the glucose time courses can be tuned by setting different PID gain parameters (Figure 8).

**Figure 8:**
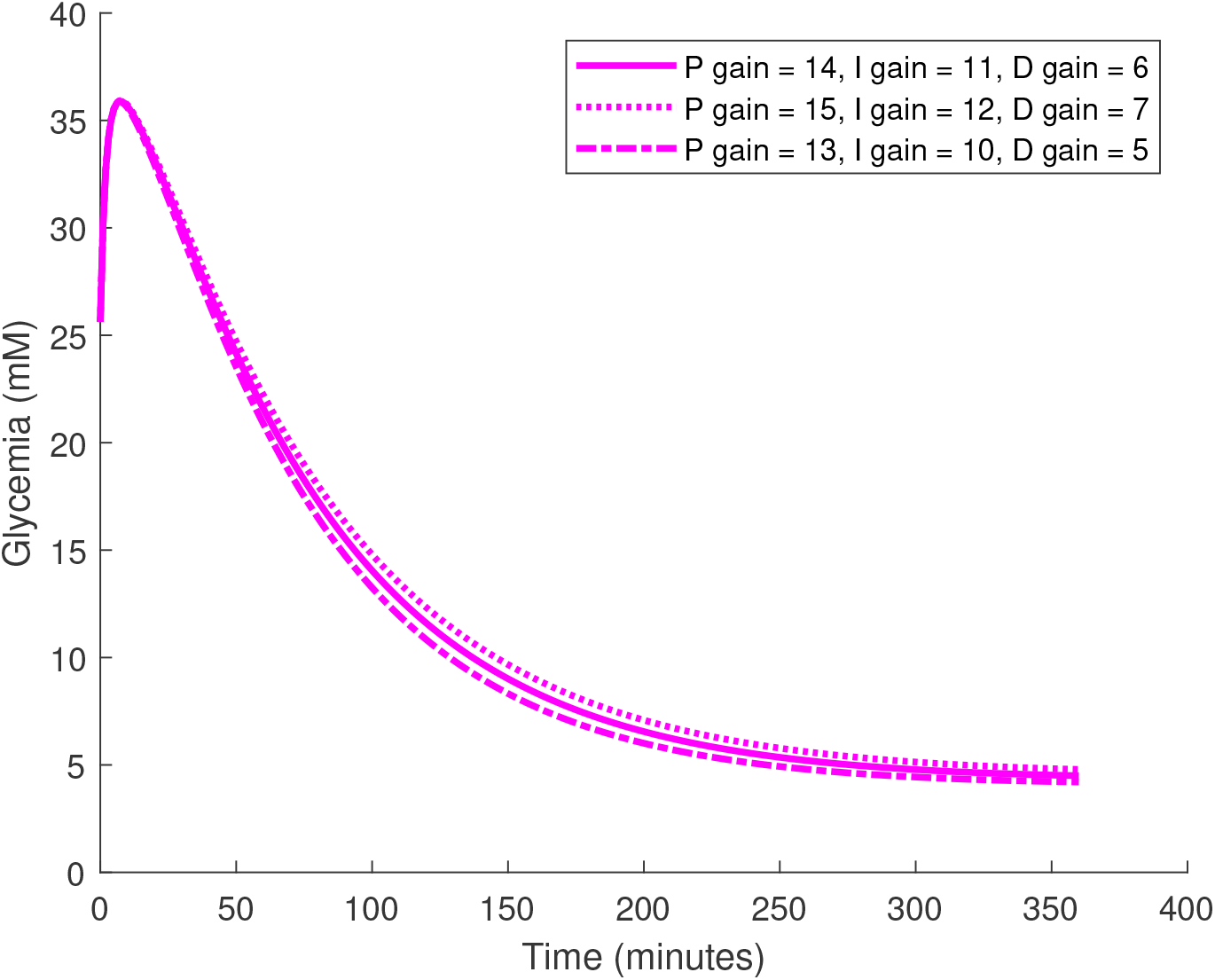
Glucose time courses for three groups of PID gain parameters. The implanted PID *β*-cell density is 5 × 10^11^ cells/L. It is noteworthy that when the PID gain parameters are increasing, the curve is actually converging more slowly, contrary to typical PID tuning patterns.

The long-term glucose level can be traded off against the glucose decrease rate. If we can tolerate a lower long-term glucose level (with the risk of hypoglycemia) we can set lower PID gain parameters for a faster decrease rate. Conversely, if we are able to tolerate a lower long-term glucose level, we may be able to accelerate the rate of glucose decrease by choosing smaller PID parameters. However, for artificial *β*-cells, the glucose curve time course cannot be changed once the cell density is fixed. We also find that the glucose time course is highly sensitive to the target value of ATP, as shown in Figure 9. In general, with a smaller ATP target value glucose decreases faster and converges to a lower level.

**Figure 9:**
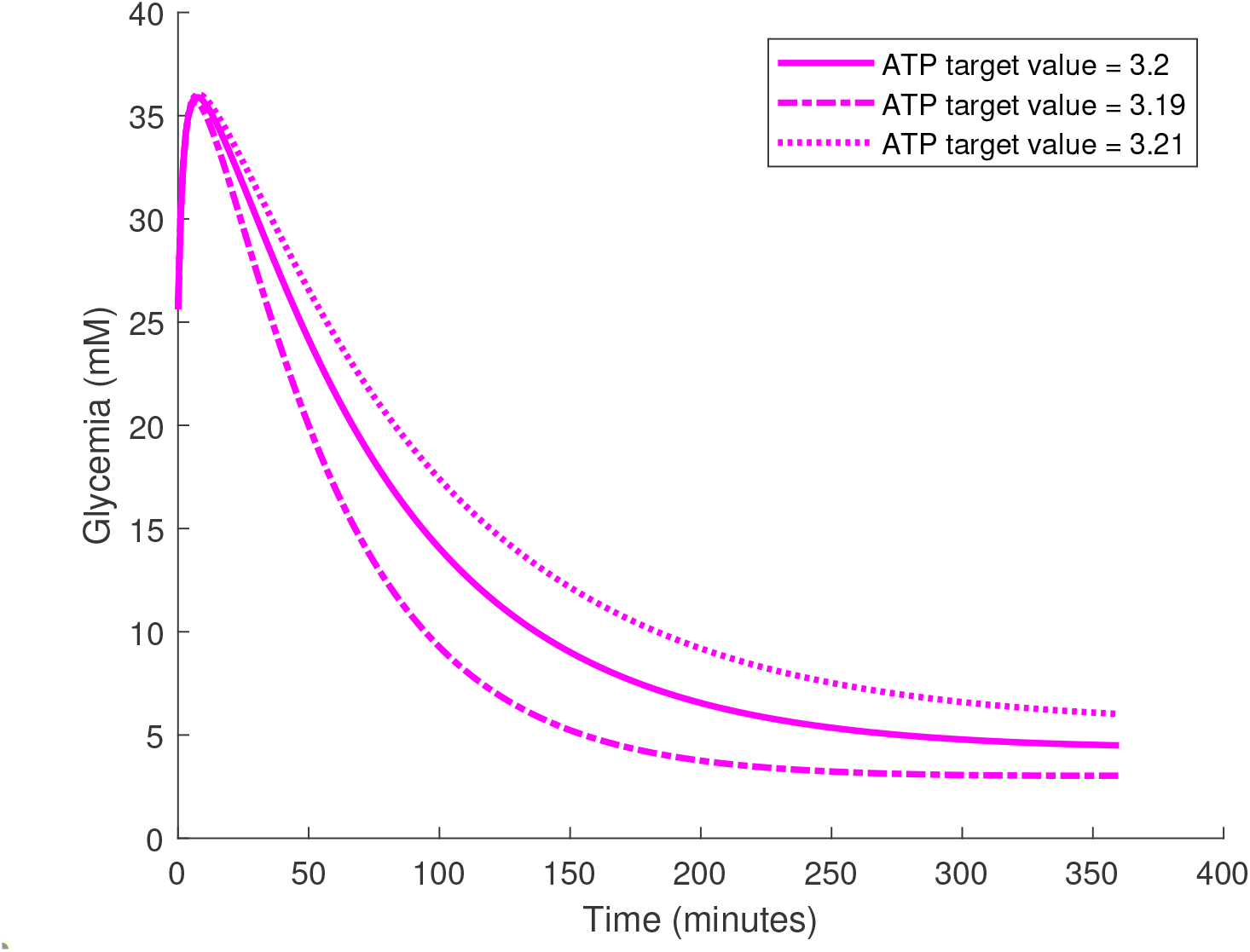
Glucose time courses as the ATP target value is varied for artificial *β*-cell. The implanted PID *β*-cell density is 5 × 10^11^ cells/L. Glucose level converges more rapidly for lower target value and vice versa.

## Discussion

By applying PID control to the *β*-cell, the glucose level is controlled with respect to the ATP level. By changing the gain parameters and the ATP target value, a PID controller is capable of providing a wide range of time course patterns and thus different glucose management outcomes. However, the final treatment result of PID *β*-cell does not reach healthy levels as quickly as healthy mice. One possible reason is that we are controlling the glucose level based on the ATP as input and mRNA as the actuator. There is a large error when the ATP is viewed as a measurement of the glucose level. Even if we can have precise reading of the glucose level, it would be difficult for the PID *β*-cell to work as well as the beta cell in healthy mice, since the beta cell in healthy mice is controlled by both the blood glucose level and the insulin level.^15^ In addition, mRNA is also a relatively weak actuator to control glucose levels, since mRNA does not directly affect glucose levels. In spite of the dramatic decrease in mRNA levels, insulin levels do not decrease as quickly as in healthy mice. PID *β*-cells can be viewed as a function with four independent variables (PID gains and ATP target values) and therefore optimization methods should be explored for parameter selection. In addition, the PID control with constant gain parameters is not optimal when controlling the glucose level. Furthermore, in nature the glucose level is controlled not just by insulin but also by glucagon, which is not provided by the reported artificial *β*-cells. The insulin production rate changes dramatically following a meal, which is a challenge for PID controllers. We plan future studies of other controller designs in search of more optimal solutions, such as variable gain parameters for PID control, fuzzy control, reinforcement learning control, and model predictive control.

## Author Information

### Data and Software Availability

The code for this paper is available in: https://github.com/liulin99/PID-beta-cell. The model does not require any additional data.

## Supplementary material

### Background on PID control

In control theory, the proportional-integral-derivative controller (PID controller) is widely used to solve the problem of maintaining a desired output level when the states are subject to frequent changes in a dynamic system. In our case, the artificial *β*-cell is a dynamic system with ion channel as the controller whose input is the ATP and whose output is mRNA. Due to the poor performance of ion channel according to the model (Figure 6), we replace it with an abstract proportional-integral-derivative controller (PID controller) that can be implemented as a chemical reaction network (CRN).

As shown in Figure 10, PID controllers are feedback control loop mechanisms widely used in applications requiring continuous modulated control. PID controllers consist of three major components: proportional operator, integral operator, and derivative operator. For an input signal *X* the proportional operator generates an output signal *k*_*p*_*X*, where the non-negative constant value *k*_*p*_ is called proportional gain. The integral operator generates an output signal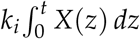, where the non-negative constant value *k*_*i*_ is called integral gain. The derivative operator generates an output signal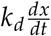, where the non-negative constant value *k*_*d*_ is called derivative gain.

**Figure 10:**
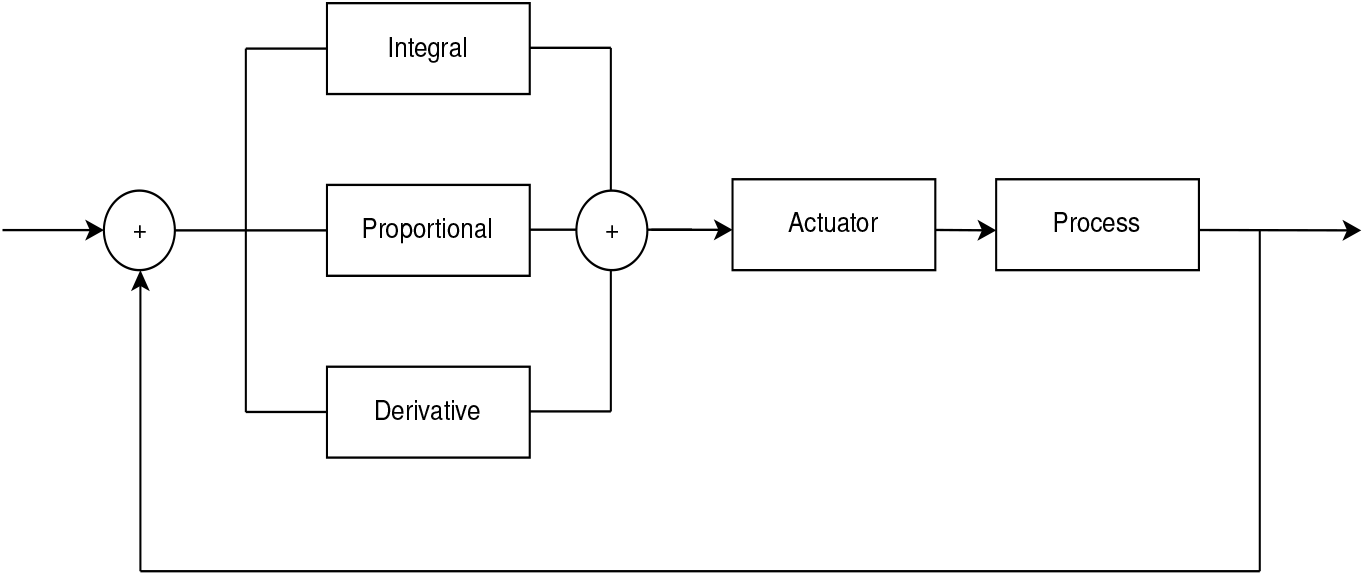
The PID-controller diagram

The actuator and the process (also known as plant) are two important concepts of a PID controller. Actuators, such as motors or heaters, generate force or energy to change the system state. Actuators may be linear or non-linear, but in either case when input to the actuator is zero, the actuator normally exerts little or no influence on the process.

An insulin-producing gene serves as an actuator, where the input is the difference between intracellular ATP concentration and a predefined target level of ATP. When this difference value is zero or negative, the expression of the insulin-producing genes is suppressed such that the secreted insulin amount is also minor. The process is the value that the actuator is trying to affect and control, the blood glucose concentration in our case. The process can be modeled as a time function *X*. In PID controllers, we have a benchmark value *X*_**benchmark**_ (also known as the set point) for the process. This is the desired target value we want our process (blood glucose level) to be. The error term, which is defined as *X* − *X*_**benchmark**_, is consistently calculated and used as the input signal to the proportional operator, integral operator, and differential operator. The output of these three components is summed and fed into the actuator to control the process. Over time, the error term changes as the process changes. The new error term as a time function is fed into the proportional operator, integral operator, and differential operator again in a feedback loop. When *X* approaches *X*_**benchmark**_, the output from the proportional operator approaches zero. Hence, the target value cannot be achieved by feeding only the output from the proportional operator into the actuator. This is called the steady state error. Integral operators can remedy the steady state error by effectively accumulating the error term and adding it to the proportional output. In this way, the actuator can maintain a higher output level even if *X* is approaching *X*_**benchmark**_. But this brings another issue: the error term is still a relatively large value even if *X* = *X*_**benchmark**_. This causes an overshoot of the process: *X* goes beyond *X*_**benchmark**_, either lower or higher, and fluctuates around *X*_**benchmark**_ for a relatively long time before it converges to *X*_**benchmark**_. The derivative operator can minimize this overshoot and let *X* converge faster to *X*_**benchmark**_.^16^

The Ziegler-Nichols tuning method is often used to experimentally find the PID controller parameters. In this method, the integral and derivative gains are first set to zero. Then we slowly increase the proportional gain until the process we are trying to control begins to oscillate. Then we gradually increase the integral and derivative gains to deal with the overshoot. However, this method fails in our setting. We tried to gradually increase the proportional gain but failed to observe any oscillation. And the process (the glucose level in our case) normally converges to the target value faster when the PID parameters are larger, which also is not observed in our case (Figure 8).

In the face of failure of standard methods we use the constrained sampling approach described in the methods and results section.

### On reproducing the model of artificial *β*-cells

We encountered the following issues when trying to reproduce the results of Xie et al.^5^

There is a term called virtual compartment in this paper. In light of the lack of details provided in the paper, it is unclear what the purpose of the virtual compartment is.

The model also has multiple tunable parameters: the initial value of glucose level, the artificial *β*-cell density, the vascular exchange constants (positive means the implanting of the cell, zero means there is no implanting), diabetic factor (zero means fully T1D mice, one means fully healthy mice). This model also involves many biology-related constants. We reconstructed this model in MATLAB and followed exactly the same protocols described in Ref. 5, however we encountered the following uncertainties and errors on some of these parameters and constants.

In equation (17) of the supplementary material of Ref. 5, a metabolic rate term *R*_4_(*Ca*_*Si*_) is defined as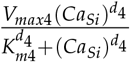. However, we fail to find the *d*_4_ value in the main paper or the supplementary material of Ref. 5. In equation (6) and equation (7) of the supplementary material of Ref. 5, a mathematical model for the synthetic compartment is described. Both equations contain a term *I*_*NaK*_(*Na*_*Si*_); however, the definition of this term is missing from Ref. 5. Equation (34) of supplementary material of Ref. 5 describes the blood insulin concentration rate of change. In this equation, there is a term *ξ*(*I*), which is defined as 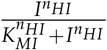 on top of page 14 of Ref. 5. The values of *K*_*MI*_ and *n*_*HI*_ are missing. Also in this equation(34), there is an *F*_*I*_ factor indicating the mice insulin production. This factor is either 0 or 1, according to Table S10 of the supplementary material of Ref. 5, where 0 means a fully diabetic mouse and 1 indicates a fully healthy mouse. But in the legend to Figure S13(D) of the supplementary material of Ref. 5, there is a variable of “native insulin production” ranging from 0.1% to 100%. We are not sure if the “native insulin production” corresponds to the *F*_*I*_ factor. And in Equation (16) of the supplementary material of Ref. 5, the unit on the right hand side does not match the left hand side. Also there is a variable *N*_*S*_ indicating the cell density value defined in Table S4 of the supplementary material of Ref. 5. This variable takes multiple values throughout the paper and the default value is 7.84 × 10^8^ cells/L as reported in Table S4 of the supplementary material of Ref. 5. In Figure 4G of the main paper of Ref. 5, there is a description of a total of 5 × 10^6^ cells implanted into the mice, but no data on the cell density is provided. We find the volume of the total capsules where the artificial *β*-cells located is 3.35 × 10^−4^ L, indicating the cell density should be 1.49 × 10^10^ cells/L. However, neither of these two density values can match what is reported in Figure 4G of the main paper of Ref. 5 via simulation. Both density values are too high (Figure 11). We contacted the authors of Ref. 5 regarding these issues, but have not received concrete answers. Hence here we have tried to correct these equations and make guesses with respect to the missing parameters consistent with the reported experimental results.

**Figure 11:**
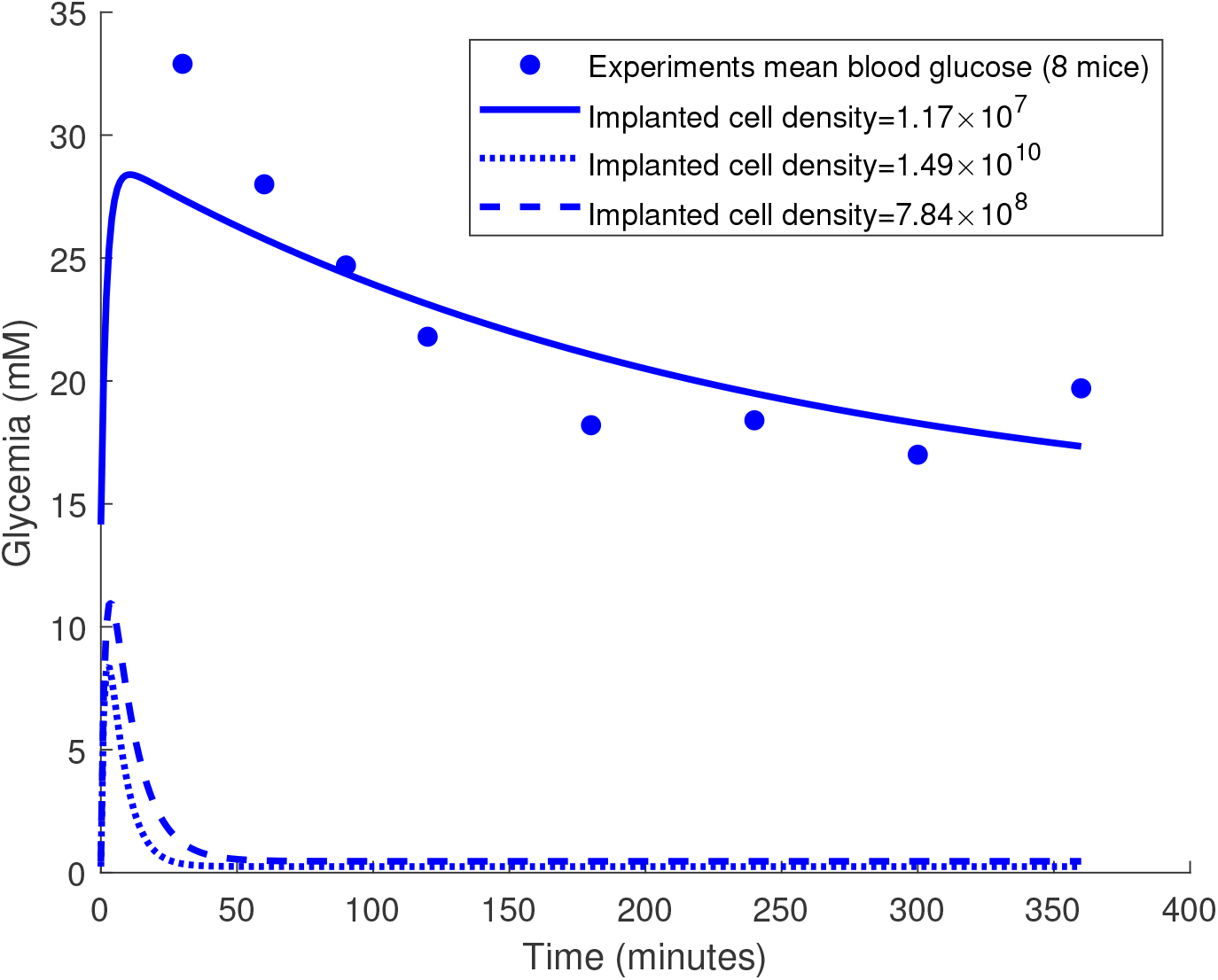
Glucose curve for different cell density values. Various values are tried and 1.17 × 10^7^ cells/L gives visually consistent results with Figure 4(G) of Xie et al. (2016). It appears that the other two density values as described in Xie et al. (2016) are too high and we fail to reconstruct their Figure 4(G) with these two values.

The following are our guesses and corrections. For the virtual compartment, we believe it to simulate the stomach, since *G*_*I*_ is described with this equation: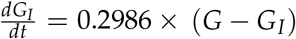, where *G*_*I*_ is the glucose concentration in the virtual compartment and *G* is the blood glucose concentration. The equation indicates in the virtual compartment, glucose is diffused into the bloodstream, just as the stomach does following a meal. There is a *d* term in Table S9 of the supplementary material of Ref. 5 with a value of 1.6. The term is described as “Hill coefficient for calcium-dependent gene expression”, which is identical to equation (17) of the supplementary material of Ref. 5. This term is not used anywhere in the paper. Our guess is the *d*_4_ parameter in equation (17) from the supplementary material of Ref. 5 is a typo and it should be *d* instead. For the *I*_*NaK*_(*Na*_*Si*_) term, we guess this is also a typo and use the clearly defined *I*_*NaK*_(*V, Na*_*Si*_) term instead. We find a same *ξ*(*I*) term from the supplementary material (page 27) of a reference^17^ cited in Xie’s paper, where *K*_*MI*_ = 10^−2^*ng*/*mL* and *n*_*HI*_ = 8. We adopted these two values in the model. We believe “1 or 0” is also a misleading description for term *F*_*I*_. We determined the correct description should be “0 to 1” and we choose this value to be 0.1% for T1D mice. As for equation(16) in the supplementary material of Ref. 5, we changed it to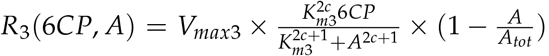, such that the dimensions are the same on both sides of this equation. For the cell density value, we tuned this parameter such that the model’s simulation results can match those reported in Figure 4G of Ref. 5 as shown in Figure 2.

### The adopted model

Equations of the artificial *β*-cell mathematical model in this paper are listed. Most of them are as in the supplementary material of Xie et al., Ref. 5. Equations that were modified, as described above, are marked with **, and the assumptions we made are marked with ***. The glucose and insulin metabolism can be described as:

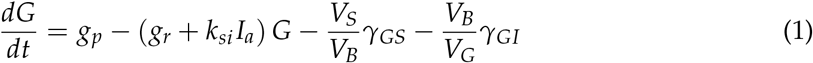

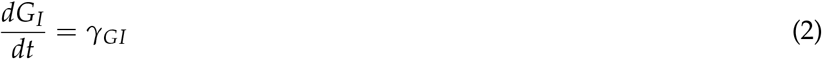

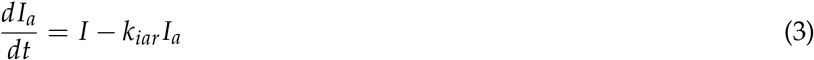

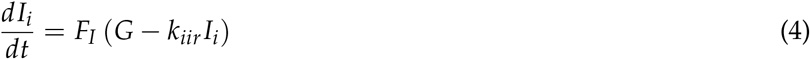

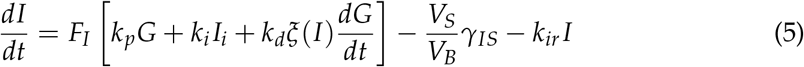

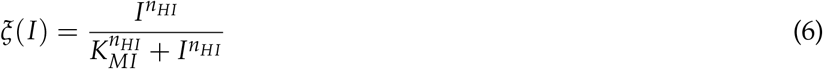

The cell metabolic rates can be described as:

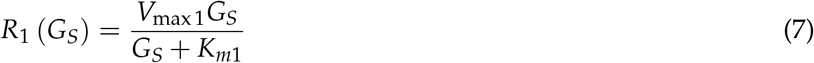

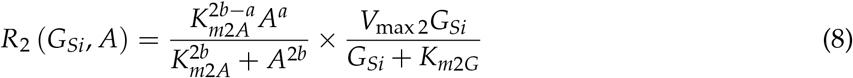

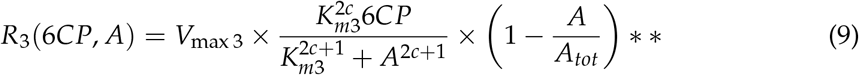

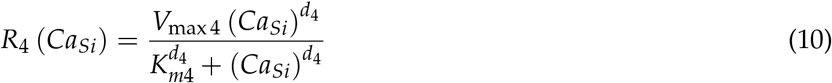

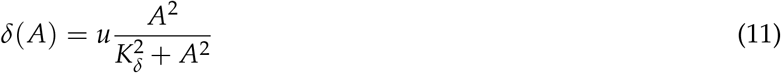

The ion channels in cell can be described as:

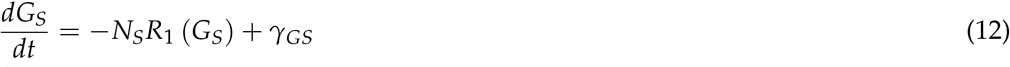

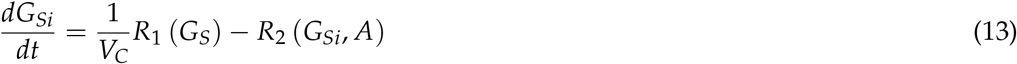

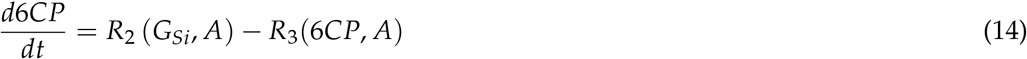

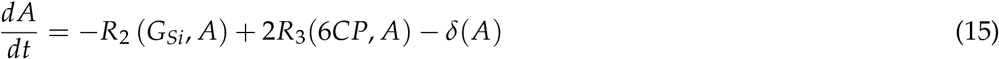

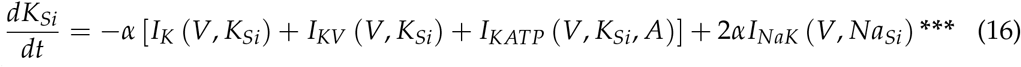

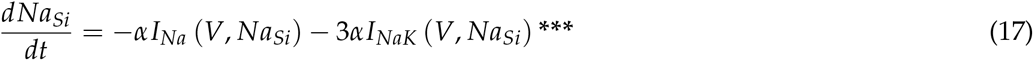

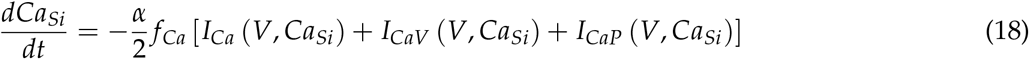

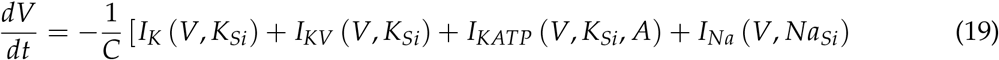

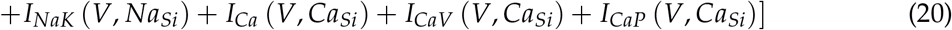

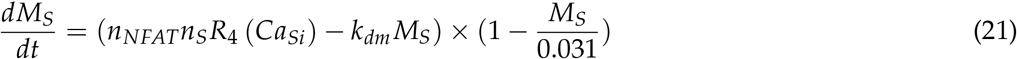

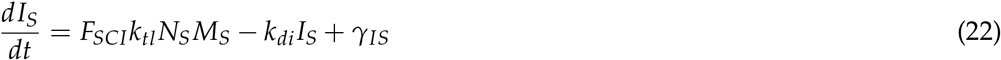

In PID *β*-cell, equation(21) is substituted by:

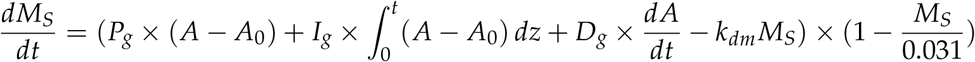

The currents and potentials in the model can be described as:

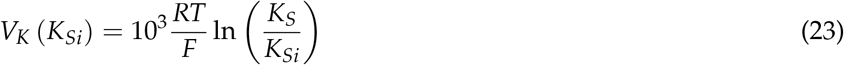

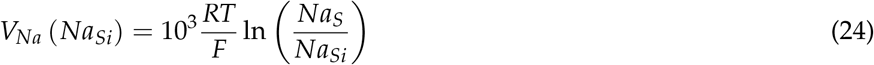

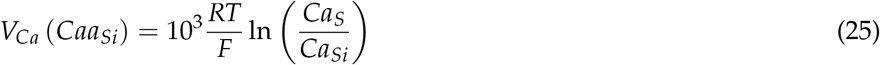

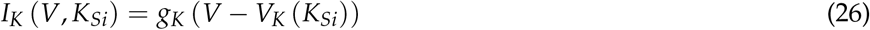

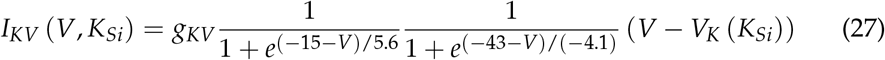

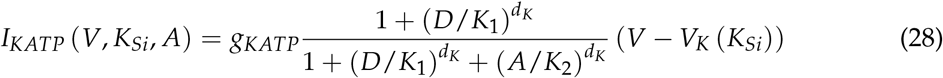

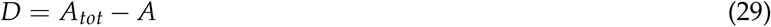

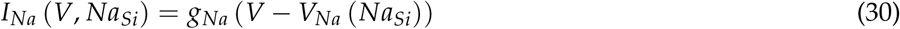

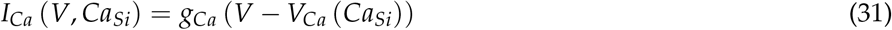

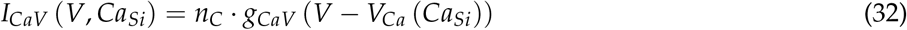

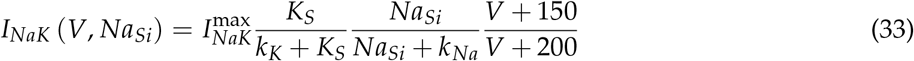

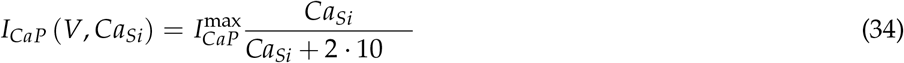

The cell flux transfer can be described as:

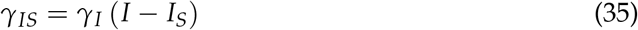

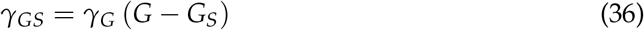

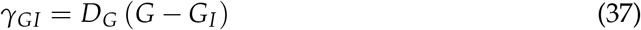

